# Follicular regulatory T cells can access the germinal centre independently of CXCR5

**DOI:** 10.1101/641589

**Authors:** Ine Vanderleyden, Sigrid C. Fra-Bido, Silvia Innocentin, Hanneke Okkenhaug, Nicola Evans-Bailey, Wim Pierson, Michelle A. Linterman

## Abstract

The germinal centre (GC) response is critical for generating high-affinity humoral immunity and immunological memory, which forms the basis of successful immunisation. Control of the GC response is thought to require follicular regulatory T (Tfr) cells, a subset of suppressive Foxp3^+^ Treg cells located within GCs. Relatively little is known about the exact role of Tfr cells within the GC, and the mechanism/s through which they exert their suppressive function. A unique feature of Tfr cells is their reported CXCR5-dependent localisation to the GC. Here we show that the lack of CXCR5 on Foxp3^+^ regulatory T cells resulted in a reduced frequency, but not an absence of, GC-localised Tfr cells. This demonstrates that additional, CXCR5-independent mechanisms facilitate Treg cell homing to the GC.

## Introduction

Follicular regulatory T (Tfr) cells are a distinct subset of Foxp3^+^ Regulatory T (Treg) cells that are located within the germinal centre (GC), where they are thought to suppress the magnitude and output of the GC response (Chung et al., 2011, Linterman et al., 2011, Wollenberg et al., 2011, Kawamoto et al., 2014, Vanderleyden et al., 2014, Sage et al., 2016, Wu et al., 2016, Botta et al., 2017, Fu et al., 2018). Tfr cells phenotypically resemble Tfh cells in many aspects, including the expression of programmed cell death protein 1 (PD-1), C-X-C chemokine receptor type 5 (CXCR5), B cell lymphoma 6 (Bcl6), Slam-associated protein (SAP), and inducible costimulator (ICOS)(Chung et al., 2011, Linterman et al., 2011, Wollenberg et al., 2011). However, Tfr cells do not express the B cell helper molecules IL-21, IL-4, and CD40L, but instead express Treg signature molecules such as GITR, CTLA-4, and Foxp3 (Chung et al., 2011, Linterman et al., 2011, Wollenberg et al., 2011, Sage et al., 2014, Wing et al., 2014). Gene expression analysis shows that Tfr cells have a distinct transcriptional profile that is more similar to Treg cells than to Tfh cells, or other T helper cell subsets (Linterman et al., 2011, Wing et al., 2017). Further, Tfr cells have suppressive function, and are therefore considered a subset of Treg cells that are thought regulate the GC response (Linterman et al., 2011, Wing et al., 2017, Stebegg et al., 2018).

Given the central role of the GC response in generating highly effective humoral immune responses and immunological memory, it is of considerable biological interest to understand how Tfr cells function within this response (Vanderleyden et al., 2014). Despite the fact that the field has grown exponentially in recent years, relatively little is known about the exact role of Tfr cells within the GC and the mechanism/s through which they exert their suppressive function. While initial studies agreed that Tfr cells can limit the size of the GC response, they lacked a system to genetically remove Tfr cells whilst leaving other Tfh and Treg cells intact (Chung et al., 2011, Linterman et al., 2011, Wollenberg et al., 2011). Therefore, we set out to develop a mouse model that specifically lacks Tfr cells, without affecting Tfh cells or other Treg cell subsets. A unique feature of Tfr cells is their location within the GC, which discriminates them from other Treg cell subsets, and this localisation was reported to be exclusively dependent on CXCR5-driven chemotaxis towards the GC (Chung et al., 2011, Wollenberg et al., 2011). In this regard, genetic removal of *Cxcr5* in Foxp3^+^ Treg cells is a logical approach for generating a mouse model that specifically lacks Tfr cells and would enable the study of the GC response in the absence of Tfr cells.

To this end, we developed three mouse strains that lack CXCR5 in all Foxp3^+^ Treg cells, or in all CD4^+^ T cells; *Cxcr5*^*fl/fl*^*Foxp3*^*cre-yfp*^, *Cxcr5*^*fl/fl*^*Foxp3*^*cre-ERT2*^ mice, and *Cxcr5*^*fl/fl*^*Cd4*^*cre/+*^ mice (Fontenot et al., 2005b, Rubtsov et al., 2010, Bradford et al., 2017). To our surprise, despite successful depletion of CXCR5 on Treg cells, Tfr cells are still present in the GC after immunisation. Loss of CXCR5 resulted in a reduced presence of Tfr cells within the GC, indicating that it is partially required for Treg cell localisation to the GC, but that is not absolutely necessary. Together, this demonstrates that CXCR5-independent mechanisms exist that allow for Treg cell localisation to the GC.

## Results

### Cxcr5^*fl/fl*^*Foxp3*^*cre*^ mice have Foxp3^+^ cells within the GC

In order to remove *Cxcr5* specifically in all Foxp3^+^ Treg cells, we crossed *Cxcr5*^*fl/fl*^ mice, in which exon 2 of *Cxcr5* was flanked by two *Loxp* sites, to *Foxp3*^*cre-yfp*^ mice (Fontenot et al., 2005b, Bradford et al., 2017). *Cxcr5*^*fl/fl*^*Foxp3*^*cre*^ mice were immunised intraperitoneally (*i.p).* with NP-KLH/Alum and the GC response in the spleen was analysed 14 days after immunisation. CXCR5 was successfully deleted in Foxp3^+^ Treg cells in *Cxcr5*^*fl/fl*^*Foxp3*^*cre*^ mice **(Figure 1A-B**). To determine whether Tfr cells were present in the GC in the absence of CXCR5, we enumerated Foxp3^+^ Treg cells present within the GC (IgD^-^Ki67^+^), by confocal imaging **(Figure 1C)**. Foxp3^+^ Tfr cells could still be identified in cryosections of the spleen of *Cxcr5*^*fl/fl*^*Foxp3*^*cre*^ mice, although their numbers were reduced by half compared with *Cxcr5*^*+/+*^*Foxp3*^*cre*^ control animals **(Figure 1D)**. Although the reduction of Tfr cells in *Cxcr5*^*fl/fl*^*Foxp3*^*cre*^ mice was modest, we hypothesised that this may result in impaired suppression of Tfh cells, and thus an increase in the number of Tfh cells. However, fewer CXCR5^+^PD-1^+^ Tfh cells were identified in *Cxcr5*^*fl/fl*^*Foxp3*^*cre*^ mice compared with controls **(Figure 1E-G)**. When Tfh cells were identified using a CXCR5-independent gating strategy based on co-expression of Bcl6 and PD-1, we observed normal frequencies and absolute numbers of Tfh cells in *Cxcr5*^*fl/fl*^*Foxp3*^*cre*^ mice **(Figure 1H-J)**. These data suggested that deletion of CXCR5 may be occurring outside the Foxp3^+^ compartment in *Cxcr5*^*fl/fl*^*Foxp3*^*cre*^ mice, as has been previously described (Franckaert et al., 2015). Indeed, a substantial population of B cells lacked CXCR5 **(Figure S1A-B)**, and deletion of the floxed exon 2 of *Cxcr5* can be detected in ear notches from *Cxcr5*^*fl/fl*^*Foxp3*^*cre*^ mice **(Figure S1C)**. Both B cells and Tfh cells use CXCR5 for migration to the GC, and, therefore, non-specific deletion of *Cxcr5* in *Cxcr5*^*fl/fl*^*Foxp3*^*cre*^ mice limits the ability to draw conclusions about the impact of the reduced frequency of Tfr cells on the GC response. Consequently, an alternative approach for deleting *Cxcr5* specifically in Foxp3^+^ Treg cells was required to determine the impact that loss of CXCR5 on Treg cells has on the GC response.

**Figure 1:**
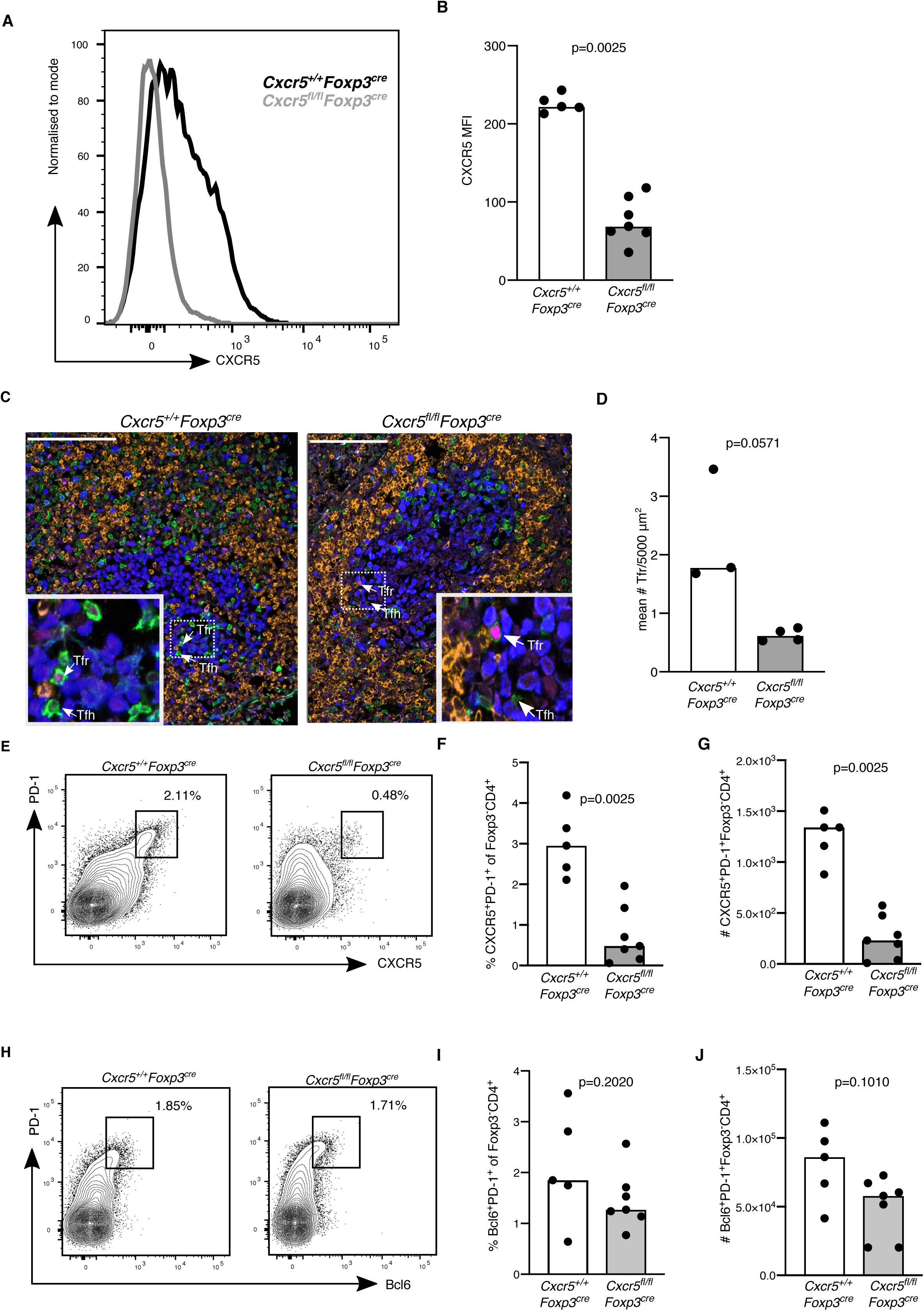
Tfr cells are present at reduced numbers in *Cxcr5*^*fl/fl*^*Foxp3*^*cre-yfp*^ mice. Mice were immunised with NP-KLH/alum *i.p.* and the germinal centre response was analysed 14 days after immunisation. **A.** Histogram of CXCR5 expression in Foxp3^+^CD4^+^ Treg cells. **B.** CXCR5 MFI (geometric mean) in Foxp3^+^CD4^+^ Treg cells. **C.** Analysis of Tfr and Tfh cells 14 days after influenza A virus (HKx31) infection in *Cxcr5*^*fl/fl*^*Foxp3*^*cre*^ mice and *Cxcr5*^*+/+*^*Foxp3*^*cre*^ controls. Representative confocal images of splenic cryosections stained for Foxp3 (magenta), Ki67 (Blue), CD3 (Green) and IgD (orange), Foxp3^+^ cells are indicated by arrows, scale bar = 40µm. **D.** Quantification of the average number of Tfr cells per mouse, defined as CD3^+^Foxp3^+^ cells within the GC, per 5000µm^2^. Each dot represents the average number of Tfr cells per 5000 µm^2^ per mouse, from 2-6 different GCs. **E.** Representative flow cytometry contour plots of CXCR5^+^PD-1^+^ Tfh cells from Foxp3^-^CD4^+^ cells. **(F-G)** Quantification of **F.** the frequency and **G.** absolute numbers of CXCR5^+^PD-1^+^ Tfh cells. **H.** Representative flow cytometry contour plots of Bcl6^+^PD-1^+^ Tfh cells of Foxp3^-^CD4^+^ cells. **(I-J)** Quantification of **I**. the frequency and **J.** absolute numbers of Bcl6^+^PD-1^+^Foxp3^-^CD4^+^ Tfh cells. Each symbol represents one mouse, and the horizontal bars represent the median values. P values were determined using a Mann-Whitney U test. Data are representative of two independent experiments.

### Specific deletion of CXCR5 in *Cxcr5*^*fl/fl*^*Foxp3*^*cre-ERT2*^ mice does not prohibit Tfr cell localisation to the GC

In an attempt to limit CXCR5 deletion specifically to Treg cells, we crossed *Cxcr5*^*fl/fl*^ mice to *Foxp3*^*cre-ERT2*^ mice, which have a cre-recombinase and mutated oestrogen receptor ligand binding domain (ERT2) inserted in the 3’ untranslated region of the *Foxp3* gene, allowing for inducible deletion of the floxed allele upon tamoxifen administration (Rubtsov et al., 2010). To induce cre-recombinase activity in Treg cells, mice received a tamoxifen containing diet for 5 weeks resulting in deletion of CXCR5 on Treg cells after 3 weeks **(Figure 2A)**. Tamoxifen treated mice were then immunised with 4-hydroxy-3-nitrophenylacetyl (NP)-Keyhole Limpet Hemocyanin (KLH/Alum) subcutaneously (*s.c.*) in the flank, followed by analysis of the inguinal LN (iLN). *Cxcr5* was successfully deleted in *Cxcr5*^*fl/fl*^*Foxp3*^*cre-ERT2*^ mice **(Figure 2 B)** without loss of CXCR5 on B cells in these animals **(Figure S2A-C).** Despite the loss of CXCR5 in Treg cells, Bcl6^+^PD-1^+^ Tfr cells could still be identified in *Cxcr5*^*fl/fl*^*Foxp3*^*cre-ERT2*^ mice, at ~40% the frequency of control mice **(Figure 2C-E)**. However, flow cytometric analysis cannot rule out that Treg cells with a Tfr cell phenotype form, but are unable to localise to the GC. Therefore, the presence of Foxp3^+^ cells within the GC were analysed in cryosections from iLN by confocal imaging **(Figure 2F).** Tfr cells could clearly be identified within the GC of *Cxcr5*^*fl/fl*^*Foxp3*^*cre-ERT2*^ mice, and quantification of the number of Tfr cells normalised to total GC area, or to the number of Tfh cells showed a 2-3-fold reduction in the number of Tfr cells within the GC, but not a complete absence of these cells **(Figure 2G-H)**. This finding is consistent with our results from the *Cxcr5*^*fl/fl*^*Foxp3*^*cre*^ mice **(Figure 1)**, but is discordant with previous reports which showed that CXCR5-deficient Treg cells did not localise to the GC after adoptive transfer into T cell deficient hosts (Chung et al., 2011). We observed that the intensity of Foxp3 expression on Treg cells within the GC is significantly lower compared with those outside of the GC, which may impair the detection of Tfr cells within the GC **(Figure 2I)**. Collectively, these data demonstrate that the deletion of CXCR5 in Treg cells is not sufficient to impair their access to the GC, suggesting that additional mechanisms to guide these cells to the GC may exist.

**Figure 2:**
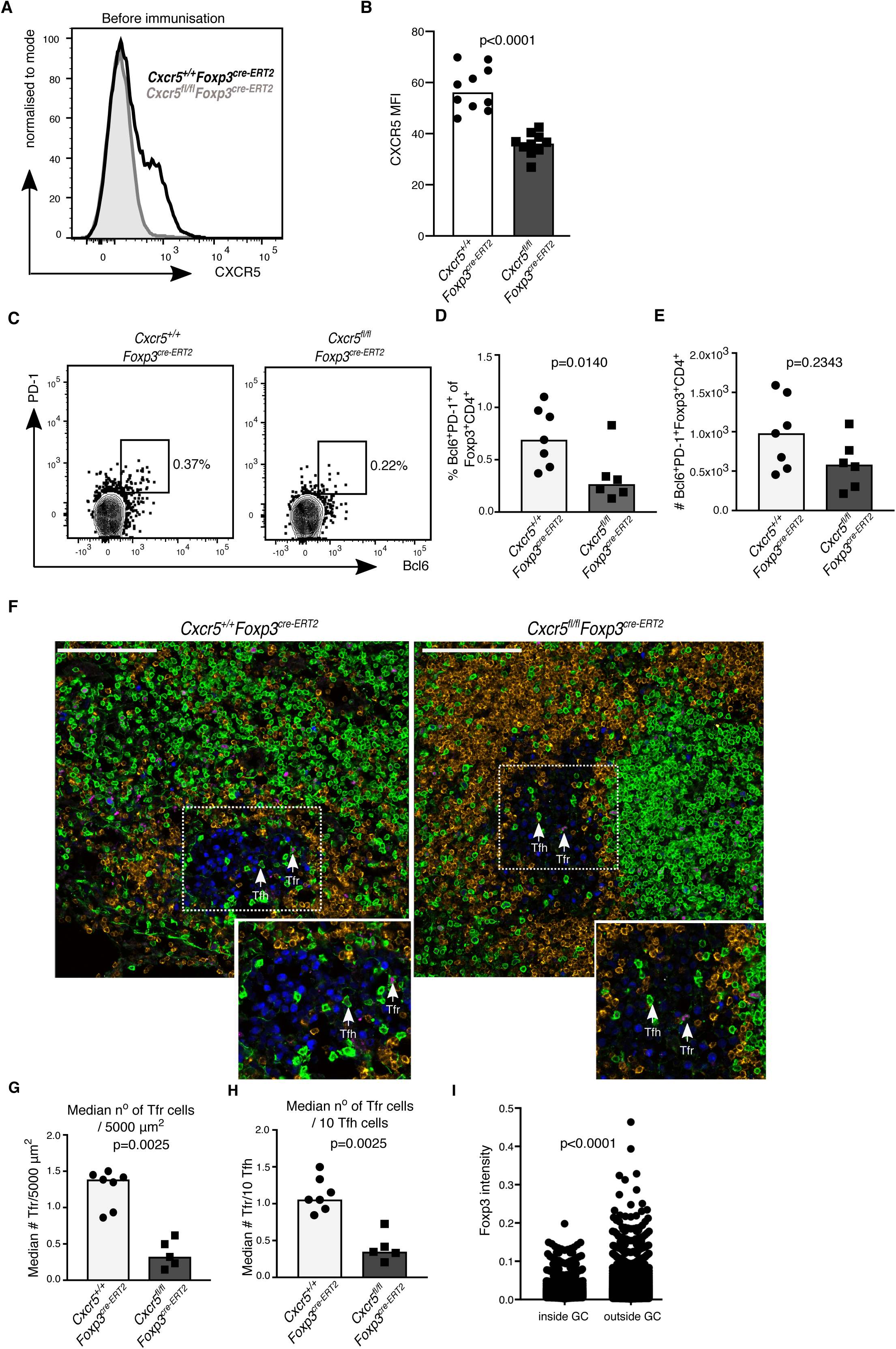
Fewer Tfr cells are present in the germinal centre of *Cxcr5*^*fl/fl*^*Foxp3*^*cre-ERT2*^ mice. Analysis of the GC response 14 days after *s.c.* immunisation with NP-KLH/Alum. **A.** Histogram of CXCR5 expression in Treg cells in mesenteric lymph nodes three weeks after initiating the tamoxifen diet, prior to immunisation, in *Cxcr5*^*fl/fl*^*Foxp3*^*cre-ERT2*^ and *Cxcr5*^*+/+*^*Foxp3*^*cre-ERT2*^ mice. **B.** Quantification of the MFI (geometric mean) of CXCR5 in Foxp3^+^CD4^+^ Treg cells. **C.** Representative flow cytometry contour plots of PD-1^+^Bcl6^+^ cells within Foxp3^+^CD4^+^ cells (Tfr cells). **(D-E)** Quantification of the **D.** frequency and **E.** absolute number of Bcl6^+^ Tfr cells. **F.** Cryosections from iLNs were stained for Foxp3 (magenta), Ki67 (Blue), CD3 (Green) and IgD (orange). scale bar = 100 µm. Representative confocal image of the GC with Tfr cells and Tfh cells indicated by the arrows. **G.** Quantification of the median number of Tfr cells, defined as CD3^+^Foxp3^+^, per 5000µm^2^. **H.** Quantification of the median number of Tfr cells per 10 Tfh cells, defined as Foxp3^-^CD3^+^. **I.** Foxp3 fluorescence intensity within and outside the GC. Each symbol represents one mouse, and the horizontal bars represent the median values. P values were determined using a Mann-Whitney U test. Data represent two independent experiments.

### The output and the size of the GC is unaltered in *Cxcr5*^*fl/fl*^*Foxp3*^*cre-ERT2*^ mice

To test whether the reduced presence of Tfr cells influenced the size and output of the GC in *Cxcr5*^*fl/fl*^*Foxp3*^*cre-ERT2*^ mice, the number of GC B cells and Tfh cells in *Cxcr5*^*fl/fl*^*Foxp3*^*cre-ERT2*^ mice were quantified. Fourteen days after immunisation the number of Ki67^+^Bcl6^+^ GC B cells was comparable in *Cxcr5*^*fl/fl*^*Foxp3*^*cre-ERT2*^ and control mice **(Figure 3A-C)**, as were PD-1^+^CXCR5^+^Foxp3^-^ Tfh cells **(Figure 3D-E).** Notably, we did not observe deletion of *Cxcr5* in conventional CD4^+^ cells (**Figure 3D**), indicating that deletion of *Cxcr5* was restricted to the Treg cell compartment in *Cxcr5*^*fl/fl*^*Foxp3*^*cre-ERT2*^ mice. To test whether the reduction in number of Tfr cells changed antibody production or affinity maturation, serum levels of NP-specific antibodies were assessed **(Figure 3G-I)**. Analysis of IgM, total IgG, and IgG1 specific for NP7 **(Figure 3G)**, or NP20 **(Figure 3H)**, or the NP20/NP7 ratio **(Figure 3I)** showed that both the titre and the quality of the humoral immune response were comparable between *Cxcr5*^*fl/fl*^*Foxp3*^*cre-ERT2*^ and control mice. Tfr cells have previously been implicated in the prevention of the outgrowth of autoreactive B cell clones within the GC (Botta et al., 2017, Fu et al., 2018). However, *Cxcr5*^*fl/fl*^*Foxp3*^*cre-ERT2*^ mice did not have elevated levels of IgG specific for double stranded DNA (dsDNA) in the serum **(Figure 3J)**, suggesting a reduction in Tfr cells does not result in a break of GC tolerance that to results in autoantibody formation after immunisation. Together, this demonstrates that loss of ~60% of the Tfr cell pool is not sufficient to alter the magnitude or output of the GC response.

**Figure 3:**
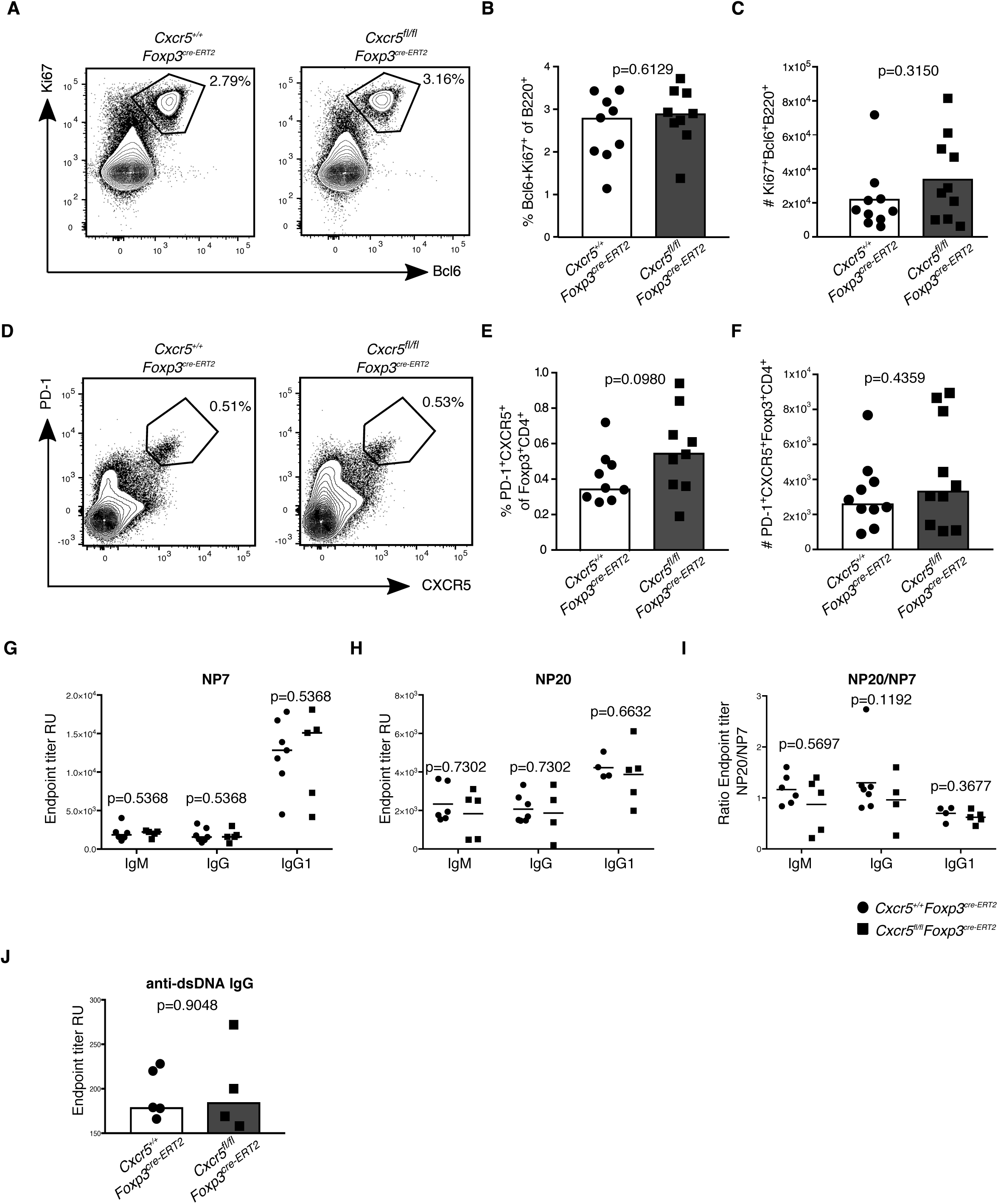
Normal magnitude and output of the germinal centre response in *Cxcr5*^*fl/fl*^*Foxp3*^*cre-ERT2*^ mice. Analysis of the GC response 14 days after *s.c.* immunisation with NP-KLH/Alum. **A.** Flow cytometry contour plots of GC B cells, gated as Bcl6^+^Ki67^+^ cells of B220^+^ cells**. (B-C)** quantification of **B.** the frequency and **C.** absolute number of Bcl6^+^Ki67^+^ B cells. **D.** Flow cytometry contour plots of Tfh cells, gated as CXCR5^+^PD-1^+^ of Foxp3^-^CD4^+^ cells. **(E-F)** Quantification of **E.** the frequency and **F.** Absolute number of CXCR5^+^ Tfh cells. **G.** Levels of IgM, IgG, and IgG1 specific for NP7 in the sera. **H.** Levels of IgM, IgG, and IgG1 specific for NP20 in the sera. **I.** Ratio of NP20/NP7 for IgM, IgG, and IgG1 in the sera. **J.** Serum levels of IgG specific for dsDNA. Each symbol represents one mouse, and the horizontal bars represent the median values. P values were determined using a Mann-Whitney U test. Data representative of two independent experiments.

### Tfh and Tfr cells are able to form in *Cxcr5*^*fl/fl*^*Cd4*^*cre/+*^ mice

The observation that Tfr cells are able to enter the GC independently of CXCR5 in both *Cxcr5*^*fl/fl*^*Foxp3*^*cre-ERT2*^ and *Cxcr5*^*fl/fl*^*Foxp3*^*cre*^ mice was unexpected as CXCR5 had previously been reported to be essential for Treg cell migration to the GC (Chung et al., 2011, Wollenberg et al., 2011). For this reason, we wished to check our observations in a third, independent, mouse model in which all T cells lack CXCR5; *Cxcr5*^*fl/fl*^*Cd4*^*cre/+*^ mice. Fourteen days after influenza A virus infection, we confirmed deletion of *Cxcr5* in Foxp3^+^ Treg cells **(Figure 4A-B)** and Foxp3^-^ Tconv cells **(Figure 4C-D)**. Despite the loss of CXCR5 in all CD4 T cells, the frequency of Ki67^+^Bcl6^+^ GC B cells was comparable to control mice with intact CXCR5 **(Figure 4E-G).** Analysis of Tfh cells and Tfr cells based on co-expression of PD-1 and Bcl6 further showed no abnormalities **(Figure S3)**. Confocal image analysis confirmed the presence of both Tfh and Tfr cells within the GC of *Cxcr5*^*fl/fl*^*Cd4*^*cre/+*^ mice **(Fig 4H-J)**, consistent with previous reports that show that CXCR5 is not essential for GC access by Foxp3^-^ CD4^+^ T cells (Moriyama et al., 2014). Taken together, these data demonstrate that lack of CXCR5 is insufficient to impair Treg cell access to the GC, suggesting that redundant mechanisms are involved in Treg cell migration to the GC.

**Figure 4:**
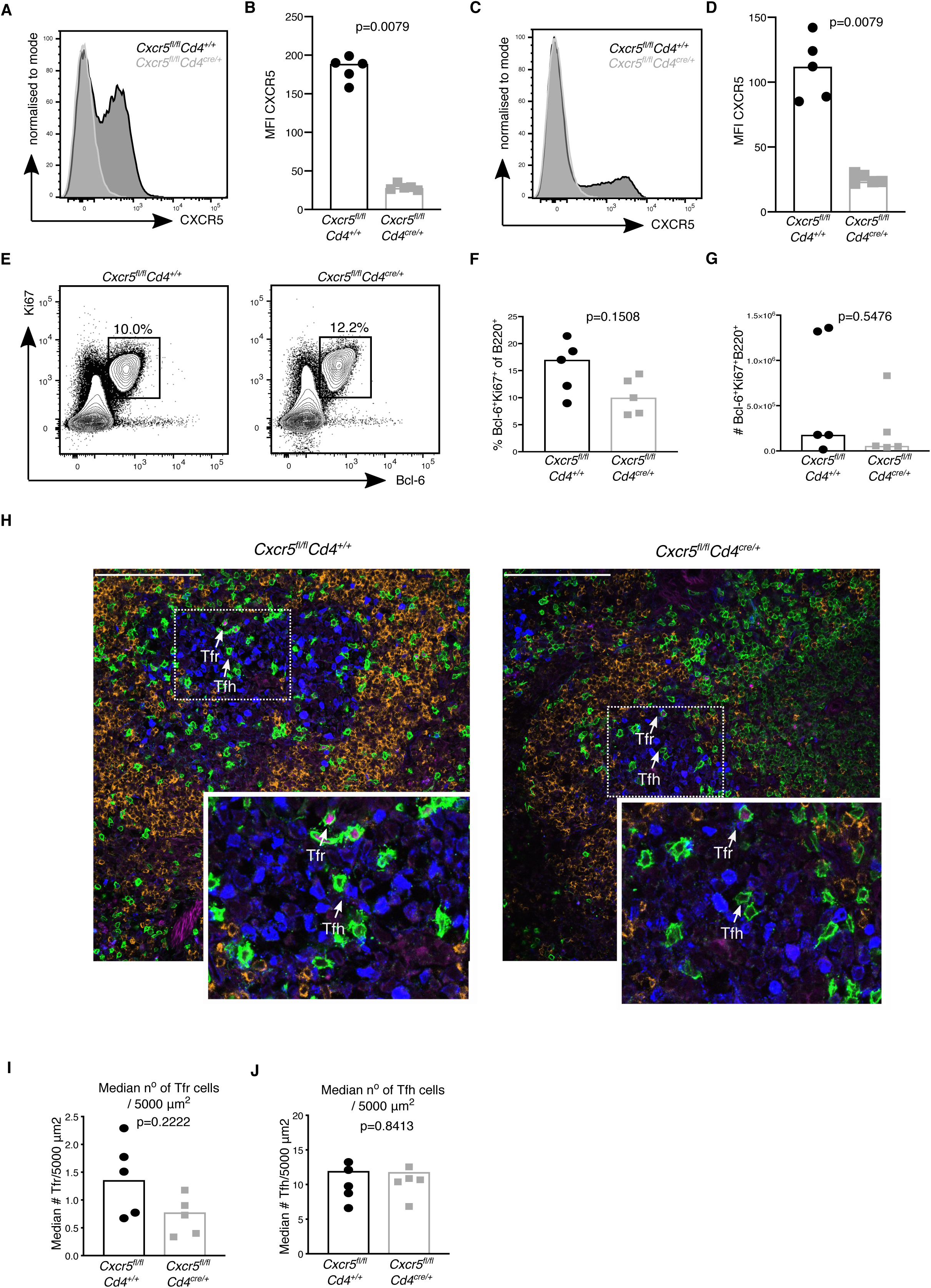
*Cxcr5*^*fl/fl*^*Cd4*^*cre/+*^ mice have an intact GC response. Analysis of the GC response in *Cxcr5*^*fl/fl*^*Cd4*^*cre/+*^ mice 14 days after influenza A virus (HKx31) infection. **A.** Representative histogram of CXCR5 expression in Foxp3^+^CD4^+^ Treg cells. **B.** Quantification of the MFI (geometric mean) of CXCR5 in Foxp3^+^CD4^+^ Treg cells. **C.** Representative histogram of CXCR5 expression in Foxp3^-^CD4^+^ T cells. **D.** Quantification of the MFI (geometric mean) of CXCR5 in Foxp3^-^CD4^+^ T cells. **E.** Flow cytometry contour plots of GC B cells, gated as Bcl6^+^Ki67^+^ cells of B220^+^ cells**. (F-G)** quantification of **F.** the frequency and **G.** absolute number of Bcl6^+^Ki67^+^ B cells. **H.** Cryosections were stained for Foxp3 (magenta), Ki67 (Blue), CD3 (Green) and IgD (orange). Scale bar = 100 µm. Representative confocal image of the GC with Tfr cells and Tfh cells indicated by the arrows. **I.** Quantification of the median number of Tfr cells, defined as CD3^+^Foxp3^+^, per 5000µm^2^. **J.** Quantification of the median number of Tfr cells, defined as CD3^+^Foxp3^+^, per 5000µm^2^. Each symbol represents one mouse, and the horizontal bars represent the median values. P values were determined using a Mann-Whitney U test. Data represent two independent experiments.

## Discussion

Tfr cells are a specialised subset of Treg cells that access the GC, where they are thought to exert suppressive functions. The localisation of Tfr cells to the GC is thought to depend on CXCR5-dependent migration to the CXCL13-rich B cell follicle. Here, we have used three independent mouse models that lack CXCR5 expression on Foxp3^+^ cells, all of which show the presence of Tfr cells within the GC. These data suggest that CXCR5 is not the only mechanism by which Tfr cells can access the GC, and that redundant mechanisms facilitate the localisation of Foxp3^+^ cells to the B cell follicle.

These were unexpected results because adoptively transferred CXCR5-deficient Treg cells into T cell-deficient mice did not migrate to the GC after immunisation (Chung et al., 2011, Wollenberg et al., 2011). In this study, we used intact mouse models rather than cell isolation and subsequent transfer, and this approach has the advantage of requiring less experimental manipulation. The differences between experimental approaches may explain the different phenotype observed. Alternatively, we noted lower expression levels of Foxp3 in Tfr cells within the GC compared with Treg cells outside the GC, consistent with their CD25^lo^ phenotype making the detection of Tfr cells more difficult than T-zone Treg cells (Wing et al., 2017). Nevertheless, the presence of CXCR5-deficient Tfr cells within the GC suggests that there are other ways by which Treg cells can access the B cell follicle.

For conventional CD4^+^Foxp3^-^ T cells to migrate to the GC, the concurrent up-regulation of CXCR5 and down-regulation of CCR7 is required to enable migration away from the T cell zone that is rich in CCR7-ligands (Haynes et al., 2007). The retention of Tfh cells within the GC is regulated by Sphingosine-1-Phosphate Receptor 2 (S1PR2), and lack of both S1PR2 and CXCR5 abrogates Tfh cell localisation to the GC (Moriyama et al., 2014). S1PR2 is also highly up-regulated in Tfr cells (Moriyama et al., 2014, Wing et al., 2017). Therefore, it is possible that S1PR2 is able to facilitate the localisation of Treg cells within the GC in *Cxcr5*^*fl/fl*^*Foxp3*^*cre-ERT2*^ mice. It is also possible that other homing receptors that are expressed on Tfr cells, such as CXCR4 whose ligand CXCL12 is expressed by the dark zone stromal cells within the GC, could be part of the redundant mechanisms involved in Tfr localisation to the GC (Denton and Linterman, 2017). The multiple mechanisms by which Treg cells migrate to different locations within tissues have likely evolved to ensure that these important suppressive cells can get to where they need to be in the absence of just one migratory cue. An understanding of the different mechanisms by which Treg cells can enter the GC may facilitate strategies to manipulate Tfr cells in both health and disease.

## Acknowledgements and funding

We acknowledge the contribution of the Babraham Institute Biological Support Unit staff, who performed *in vivo* treatments of our animals and took care of animal husbandry. We thank the staff of the Babraham Flow Cytometry Facility and Imaging Facility for their technical support. This study was supported by funding from the Biotechnology and Biological Sciences Research Council (BBS/E/B/000C0407, BBS/E/B/000C0427 and the Campus Capability Core Grant to the Babraham Institute), the European Research Council (637801 TWILIGHT). I. Vanderleyden was funded by a Cambridge Trust European Scholarship.

## Author Contributions

Authors contributions: I. Vanderleyden designed the study, performed experiments, analysed results, and wrote the manuscript. S.C. Fra-Bido and S. Innocentin performed experiments. H Okkenhaug assisted with image analysis. W. Pierson and N. Evans-Bailey provided scientific advice and help with experimental design. M. A Linterman designed and supervised the study, obtained funding and wrote the manuscript. All authors read, edited, and approved the manuscript.

## Declaration of Interests

The authors declare no competing financial interests.

**Figure S1: Non-specific deletion of CXCR5 in *Cxcr5***^***fl/fl***^***Foxp3***^***cre***^ **mice. A.** Representative flow cytometry histogram. **B.** Quantification of CXCR5 expression on B220^+^ B cells in mice of the indicated genotype. **C.** Image from gel electrophoresis showing results from a PCR reaction for the wild type (Wt) *Cxcr5* allele (214bp), *Cxcr5* excised allele (292bp) and the *Cxcr5* floxed allele (375bp) using ear notches from *Cxcr5*^*fl/fl*^*Foxp3*^*cre*^ mice, and *Cxcr5*^*fl/fl*^*Foxp3*^*+(/+)*^ and *Cxcr5*^*+/fl*^*Foxp3*^*+/+*^ controls. Each symbol represents one mouse, and the horizontal bars represent the median values. P values were determined using a Mann-Whitney U test. Data represent two independent experiments.

**Figure S2: *Cxcr5***^***fl/fl***^***Foxp3***^***cre-ERT2***^ **mice show specific deletion of CXCR5 in Foxp3**^**+**^ **Treg cells.** Analysis of CXCR5 expression on B cells 14 days after *s.c*. immunisation with NP-KLH/Alum. **A.** Representative flow cytometry contour plots of B220^+^ B cells from single cells. **(B-C)** Quantification of **B.** the frequency and **C.** absolute numbers of CXCR5^+^ B220^+^ cells. Each symbol represents one mouse, and the horizontal bars represent the median values. P values were determined using a Mann-Whitney U test. Data represent three independent experiments.

**Figure S3: Bcl6**^**+**^ **Tfh and Tfr cells form in *Cxcr5***^***fl/fl***^***Cd4***^***cre/+***^ **mice.** Analysis of the GC response in *Cxcr5*^*fl/fl*^*Cd4*^*cre/+*^ mice 14 days after influenza A virus (HK/x31) infection **A.** Flow cytometry contour plots of Tfh cells, gated as Bcl6^+^PD-1^+^ cells within CD4^+^ Foxp3^-^ cells**. (B-C)** quantification of **B.** the frequency and **C.** absolute number of Bcl6^+^PD-1^+^ Tfh cells. **D.** Flow cytometry contour plots of Tfh cells, gated as Bcl6^+^PD-1^+^ cells within CD4^+^ Foxp3^-^ cells**. (E-F)** quantification of **E.** the frequency and **F.** absolute number of Bcl6^+^ Tfh cells. Each symbol represents one mouse, and the horizontal bars represent the median values. P values were determined using a Mann-Whitney U test. Data represent two independent experiments.

## Methods

### Mice

The following mice were used in this study: *Cxcr5*^*fl/fl*^ (Bradford et al., 2017), *Foxp3*^*cre-yfp*^ (Fontenot et al., 2005a), *Foxp3*^*EGFP-cre-ERT2*^ (Rubtsov et al., 2010), *Cd4*^*cre*^ (Lee et al., 2001), *Cxcr5*^*fl/fl*^ *Foxp3*^*cre-YFP*^, *Cxcr5*^*fl/fl*^ *Foxp3*^*EGFP-cre-ERT2*^, and *Cxcr5*^*fl/f*^ *Cd4*^*cre*^ mice. All mice were on the C57BL/6 background. Mice were between three and 12 weeks old at the start of the experiment, and age- and sex-matched controls were used, unless stated otherwise. Mice were bred and maintained in the Babraham Institute Biological Support Unit. No primary pathogens or additional agents listed in the FELASA recommendations were detected during health monitoring surveys of the stock holding rooms. Ambient temperature was ~19-21°C and relative humidity 52%. Lighting was provided on a 12 hour light: 12 hour dark cycle including 15 min ‘dawn’ and ‘dusk’ periods of subdued lighting. After weaning, mice were transferred to individually ventilated cages (GM 500: Techniplast) with 1-5 mice per cage. Mice were fed CRM (P) VP diet (Special Diet Services) ad libitum and received seeds (e.g. sunflower, millet) at the time of cage-cleaning as part of their environmental enrichment. All mouse experimentation was approved by the Babraham Institute Animal Welfare and Ethical Review Body. Animal husbandry and experimentation complied with existing European Union and United Kingdom Home Office legislation and local standards.

### Immunisation and Influenza Infection

NP-KLH (Conjugation ratio 29-33, Biosearch Technologies) was dissolved in PBS to 1 mg/ml and mixed with Imject Alum (ThermoFisher Scientific) in a 1:1 ratio by vortexing to a final working concentration of 0.5 mg/ml. Mice were immunised *i.p.*, or *s.c*. on each side of the hind flank with 100µl per flank of NP-KLH/Alum, and inguinal LN were harvested 14 days after immunisation. Blood was collected after euthanasia in each experiment by cardiac puncture to determine NP-specific antibody production. For influenza infection, mice were inoculated intranasally (*i.n.*) with 10^4^ plaque-forming units of the influenza A/HK/x31 (H3N2) virus under inhalation anaesthesia using isoflurane. The mediastinal LN was harvested 14 dpi.

### Tamoxifen Treatment

Inducible deletion of floxed alleles mediated by the cre-recombinase linked to a human mutated oestrogen ligand binding receptor (ERT2) was achieved by supplementing the food with tamoxifen. From the age of three weeks, mice received a soy-free, 16 % global protein rodent diet (Teklad) for ten days. After ten days, mice were fed *ad libitum* up until six weeks with Tecklad CRD TAM^400/CreER^ Tamoxifen pellets (Teklad), containing 400mg tamoxifen citrate/kg (w/v), softened in 20% (w/v) sucrose in water solution (Sigma).

### Flow cytometry

A single cell suspension was prepared by pressing the LN through a 40-um cell strainer in 2% foetal bovine serum in PBS before antibody staining in Brilliant Stain buffer (Becton Dickinson). Antibodies used were as follows: B220 (RA3-6B2, Biolegend), Bcl6 (K112-91, BD), CD4 (RM4-5, Biolegend), CD4 (GK1.5, ThermoFisher Scientific), CD44 (IM7, Biolegend), CXCR5 (L138D7, Biolegend), Foxp3 (FJK-16S, ThermoFisher Scientific), Ki67 (SolA15, ThermoFisher Scientific), PD-1 (RMP1-30). Cells were fixed and permeabilised for intracellular staining using the eBioscience Foxp3/ Transcription Factor Fixation/Permeabilisation Staining buffer set (ThermoFisher Scientific) according to manufacturer’s instructions. Dead cells were excluded by using the zombie aqua fixable viability dye (Biolegend).

### Immunofluorescence imaging

Preparation of frozen LN samples and immunofluorescence staining was performed as described previously (Vanderleyden and Linterman, 2017). Images were acquired with a Zeiss 780 microscope using 20x and 40x objectives. Image analysis was performed using Volocity (PerkinElmer).

### ELISA

For the NP-specific ELISA, Nunc Maxisorb 96-well plates (ThermoFisher Scientific) were coated with NP7-BSA (Biosearch Technologies) at 10µg/ml or NP20-BSA (Biosearch Technologies) at 2.5µg/ml, and incubated overnight at 4°C. To determine serum levels of IgG specific for dsDNA, Nunc Maxisorb 96-well plates were coated with 100µl poly-l-lysine H_2_0 at 20 µg/ml (Sigma, cat # P4832) overnight at 4°C. Serum samples were serially diluted, and Horseradish Peroxidase (HRP) conjugated goat anti-mouse IgG1 (Abcam), IgM (Abcam) or IgG (Abcam) were added. Plates were developed using the 3,3’,5,5’-Tetramethylbenzidine (TMB) substrate set (Biolegend). Plates were read at 450nm using a PHERAstar FS plate reader (BMG Labtech).

### Statistical analysis

Statistical tests were chosen in advance as part of the experimental design. Sample sizes were determined in advance based on the availability of age-matched experimental mice and controls. Unpaired comparisons were performed using the Mann-Whitney U test. All data points were analysed and outliers were not removed unless there were technical errors. Data are presented as the median with single data points. p< 0.05 was used as a threshold for statistical significance.

