## Supplementary figures and images for "Follicular regulatory T cells can access the germinal centre independently of CXCR5"

### Supplementary figures

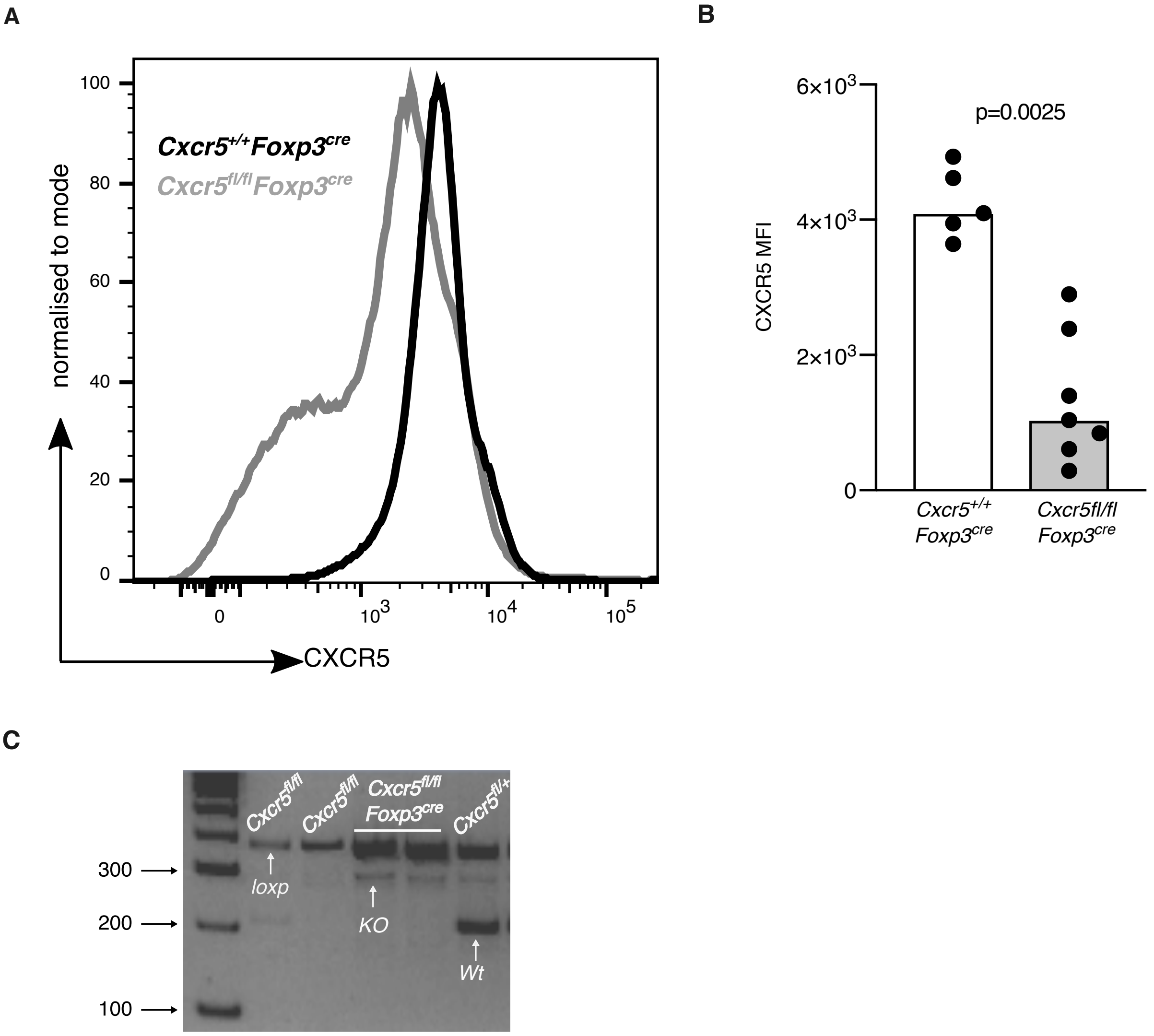

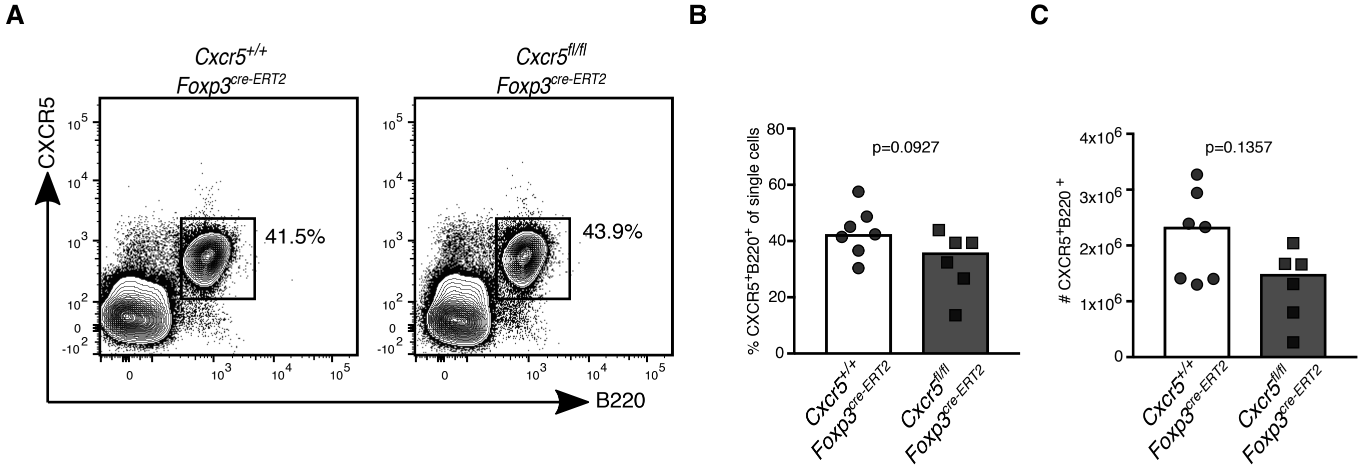

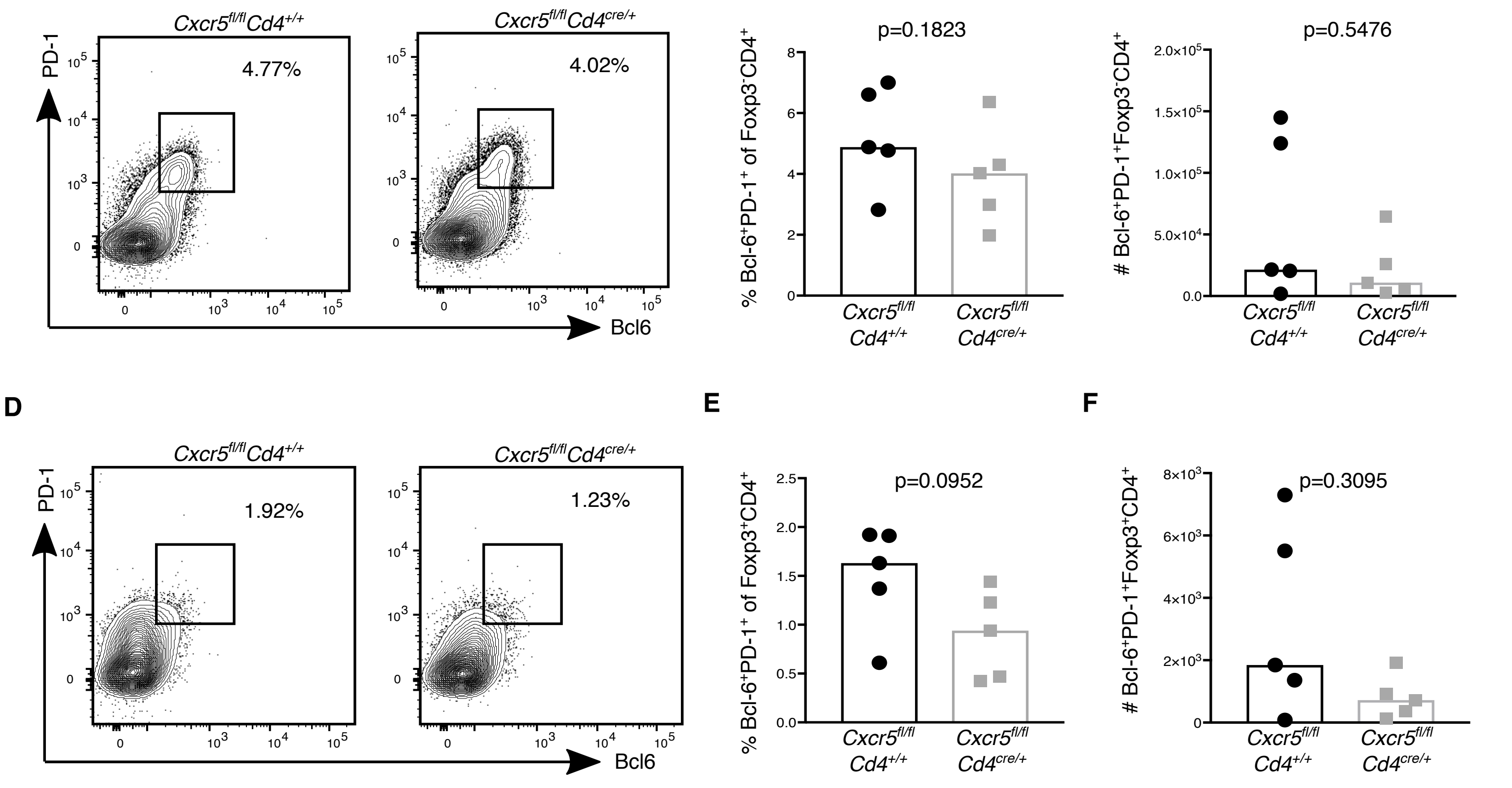
